# The overdue promise of short tandem repeat variation for heritability

**DOI:** 10.1101/006387

**Authors:** Maximilian O. Press, Keisha D. Carlson, Christine Queitsch

**Author notes:** Corresponding author: Queitsch, C.

## Abstract

Short tandem repeat (STR) variation has been proposed as a major explanatory factor in the heritability of complex traits in humans and model organisms. However, we still struggle to incorporate STR variation into genotype-phenotype maps. Here, we review the promise of STRs in contributing to complex trait heritability, and highlight the challenges that STRs pose due to their repetitive nature. We argue that STR variants are more likely than single nucleotide variants to have epistatic interactions, reiterate the need for targeted assays to accurately genotype STRs, and call for more appropriate statistical methods in detecting STR-phenotype associations. Lastly, somatic STR variation within individuals may serve as a read-out of disease susceptibility, and is thus potentially a valuable covariate for future association studies.

## The ‘missing heritability’ of complex diseases and STR variation

Complex diseases such as diabetes, various cancers, cardiovascular disease, and neurological disorders cluster in families, and are thus considered to have a genetic component [1–3] (Glossary). The identification of these genetic factors has proven challenging; although genome-wide association (GWA) studies have identified many genetic variants that are associated with complex diseases, these generally confer less disease risk than expected from empirical estimates of heritability. This discrepancy, termed the ‘missing heritability’, has been attributed to many factors [1–6]. A trivial explanation is that shared environments among relatives may artificially inflate estimates of heritability. However, missing heritability may also be due to variants in the human genome that are currently inaccessible at a population scale [1,2]. One such class of variation is short tandem repeat (STR) unit number variation. Some have previously suggested that adding STR variation to existing genetic models would considerably increase the proportion of heritability explained by genetic factors in human disease [7,8]. Three percent of the human genome consists of STRs [9] and 6% of human coding regions are estimated to contain STR variation [10,11]. Recently, the first catalog of genome-wide population-scale human STR variation has appeared [12], opening up new possibilities for understanding the contribution of STRs to human genetic diseases. This catalog, and similar data sources [13], have appeared decades after initial calls for the assessment of the role of STRs in phenotypic variation [14], lagging behind surveys of other genomic elements. Much of the initial interest in STRs was generated by the discovery of phenomena such as genetic anticipation, which are mediated by the unique features of STRs [15]. As we will discuss, new and forthcoming data sources will help to realize the long-deferred promise of STRs for explaining heritability.

STRs consist of short (2-10 bp) DNA sequences (units) that are repeated head-to-tail multiple times. This structure causes frequent errors in recombination and replication that add or subtract units, leading to STR mutation rates that are 10-fold to 10^4^-fold higher than those of non-repetitive loci [16,17]. Due to technical barriers, STR variation has until very recently remained inaccessible to genome-wide assessment.

STRs are often conserved (even if their unit number or even sequence changes), especially in coding sequences [18–21]. In both humans and the yeast *Saccharomyces cerevisiae*, promoter regions are known to be dramatically enriched for STRs [22,23]. In coding regions, STRs tend to occur in genes with roles in transcriptional regulation, DNA binding, protein-protein binding, and developmental processes [16,21,22]. These consistent functional enrichments across vastly diverged lineages suggest important functional roles for STRs.

Indeed, analysis of STR variation in the Drosophila Genetic Reference Panel identified dozens of associations between STR variants and quantitative phenotypes in recombinant inbred fly lines [13]. Moreover, accumulating evidence from exhaustive genetic studies shows that STR variation has dramatic, often background-dependent phenotypic effects in model organisms [25–29]. Together, these findings suggest that STR variation has the potential to dramatically revise the heritability estimates attributable to genetic factors.

The high STR mutation rate also leads to substantial somatic variation of STR loci within individuals. In fact, this somatic variation, also called microsatellite instability (MSI), has been used for decades as a biomarker for different classes of cancer [30]. Recent studies demonstrate that organisms exposed to various environmental stresses and perturbations show increased genome instability, including MSI [31–34]. MSI may be useful as a biomarker for cellular stress states that may predispose to disease.

The broad interest in STR variation has led to the development of techniques for high-throughput genotyping of STRs [35,36] and an explosion of analysis tools for extracting STR variation from existing sequence data [37–39]. However, the precision of these methods remains limited, due to a combination of low effective coverage of STRs and the lack of robust models for distinguishing technical error from somatic variation. Attempts to use STR variation for GWA in a fashion equivalent to SNV variation may be underpowered and confounded by the unique characteristics of this class of variants. In this review, we discuss the latest advances in these fields, and lay out a set of priorities for the future study of STRs.

## STR variation is associated with human genetic diseases

Within coding regions, STR mutations are generally in-frame additions and subtractions of repeat units, resulting in proteins with variable, low-complexity amino acid runs [21]. These mutations can result in phenotypic effects and lead to genetic disorders; several neurological diseases (spinocerebellar ataxias, Huntington’s disease, spinobulbar muscular atrophy, dentatorubral-pallidoluysian atrophy, intellectual disability, etc.) are a consequence of dramatically expanded STR alleles [7,40,41]. Many of these disease-associated STR expansions behave as dominant gain-of-function mutations [7]. However, even comparatively modest coding STR variation may confer disease risk or behavioral phenotypes, according to a variety of single-marker association studies [42–45]; for instance, variants in separate coding STRs in *RUNX2* are associated with defects in bone mineralization, higher incidence of fractures [46,47]; STR variation in this gene in dogs is also associated with craniofacial phenotypes [48]. Noncoding STR variation in regulatory sequences can affect transcription, RNA stability, and chromatin organization. For instance, certain STR variants alter *CFTR* expression and thus cystic fibrosis status [16]. We take these studies as evidence that STR variation, even in the absence of large expansions, may contribute significantly to the heritability of human traits and genetic diseases.

The severity of the STR expansion-associated diseases may suggest that natural selection should eliminate STRs in functional regions, but several recent studies across many organisms indicate that variable STRs are globally maintained [19,20,24,49,50]. For example, the pre-expansion polyQ-encoding STR in the human gene SCA2 is under positive selection, suggesting that this variable STR is actively maintained in spite of the pathogenic expansions that do occasionally occur and cause spinocerebellar ataxia [51]. Considering both the evidence of positive selection on STRs and the functional enrichments of STR-containing genes, several authors have proposed that functional STRs are maintained because they confer ‘evolvability’, or the capacity for fast adaptation [21,22,52–54]. This suggestion is intriguing, in part because many STR mutations are dominant, and, when beneficial, can quickly sweep to fixation. Although we do not further discuss these evolutionary considerations here, they underscore the phenotypic potential of STR variation.

## STR variation has dramatic background-dependent effects on phenotype

To date, the functional consequences of unit number variation in selected STRs have been studied in plants, fungi, flies, voles, dogs, and fish [25,27,28,55–57], among other organisms. In *Saccharomyces cerevisiae*, STR unit number in the *FLO1* gene accurately predicts the phenotype of cell-cell and cell-substrate adhesion (flocculation); flocculation provides protection against various stresses [57,58]. STR variation in yeast promoters has been shown to alter gene expression [22]. In *Drosophila melanogaster*, *Neurospora crassa*, and *Arabidopsis thaliana*, natural coding STR variation in circadian clock genes alters diurnal rhythmicity and developmental timing [25–27,59]. Some have proposed that the large phenotypic responses to selection observed in the Canidae are a consequence of elevated STR mutation rates relative to other mammalian clades [48,53]. We can state unambiguously that naturally variable STRs underlie dramatic phenotypic variation in model organisms.

Beyond the observable fact that variable STRs affect phenotype, we can make specific predictions about the components of phenotypic variation that they affect. Both theoretical expectations and empirical data indicate that STR variants are likely to participate in epistatic interactions, and probably more so than most SNVs. One plausible hypothesis is that STRs act as mutational modifiers of other loci, as may be expected intuitively from their elevated mutation rate (Box 1, Figure I).

### Box 1

#### Modifier mutations leading to epistasis are expected in STRs.

We have previously proposed that STRs might be more susceptible to genetic interactions [25], as we will briefly explicate here. Consider a simple two-locus haploid model under panmixis, in which loci *A* and *B* each start with a single allele (*ab*) and have the same probability *p* per generation of mutating to a second allele (*a\** or *b\**), with *p* also as the probability per generation of reverting mutations (Figure I). Let us further assume that *A* and *B* are in sign epistasis [99] (that is, *a*b* and/or *ab\** have fitness less than *ab* and *a*b\**). To escape the unfavorable *a*b* genotype, the organism may either revert to *ab* or mutate forward to *a*b\**. When the *A* and *B* loci have equal mutation rates, we expect that the reversion of a single mutant is just as likely as a second mutation, and consequently that *a*b\** individuals will appear only relatively rarely and slowly. However, consider a similar model, in which locus B has an elevated mutation rate *p_b_* > *p_a_*. In this case, the *a*b* genotype has a higher probability of a second, modifying mutation to *a*b\** than of a reversion to *ab*. Moreover, flux along the other mutational path (*ab* → *ab\** → *a*b\**) will be increased. In sum, *a*b\** genotypes will arise at higher rates, and will attain their equilibrium frequency much more rapidly, if either A or B has an elevated mutation rate [100] (p.131). This scenario can lead quickly to an equilibrium population in which incompatible epistatic alleles are frequent, even though recombinants have lower fitness. Relaxing the assumption of no population structure will further speed this process. Consequently, we would expect STRs and other loci with high mutation rates to be more likely to modify other alleles than loci with lower mutation rates, as long as we assume that all loci are equally capable of genetic interactions. This process may be referred to as ‘coadaptation’. For a rigorous model of the evolution of hybrid incompatibility, see Orr [101].

**Figure I.**
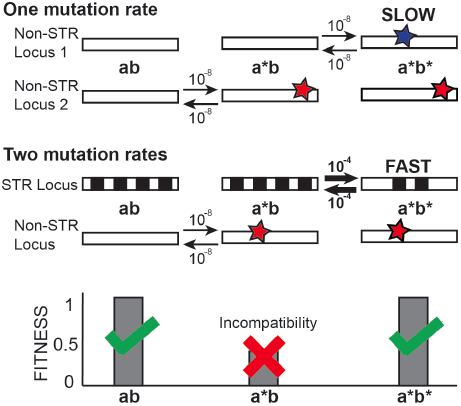
A locus with higher mutation rates allows genetic modification of unfavorable genotypes at interacting loci. Top, a model of evolution under epistasis with only one slow mutation rate. Middle, a model of evolution under epistasis with a slow and a fast mutation rate. Boxes represent loci, stars represent SNV-type mutations, black and white checkering indicates an STR locus (*a/b*, *a*/b*, and *a*/b\** signify different genotypes). Arrows with numbers represent possible mutations and their respective rates. Bottom, fitness of each genotype under both models. We expect that the model with two mutation rates will occupy the fully derived state (*a*/b\**) more quickly.

This expectation is borne out in the handful of studies reporting exhaustive genetic analysis of STRs. For instance, in the *Xiphophorus* genus of fish, a genetic incompatibility has recently been attributed to the interaction between the *xmrk* oncogene and an STR in the promoter of the tumor suppressor *cdkn2a/b* [29,60]. If the *xmrk* gene product is not properly regulated by *cdkn2a/b*, fish develop fatal melanomas, a two-locus Bateson-Dobzhansky-Muller incompatibility described in classic genetic experiments (Figure 1A) [61–63]. Expansions in the *cdkn2a/b* promoter STR are associated with the presence of a functional copy of the *xmrk* oncogene across species, and are thought to functionally repress the activity of the *xmrk* gene product through increased dosage of the tumor suppressor [29].

**Figure 1.**
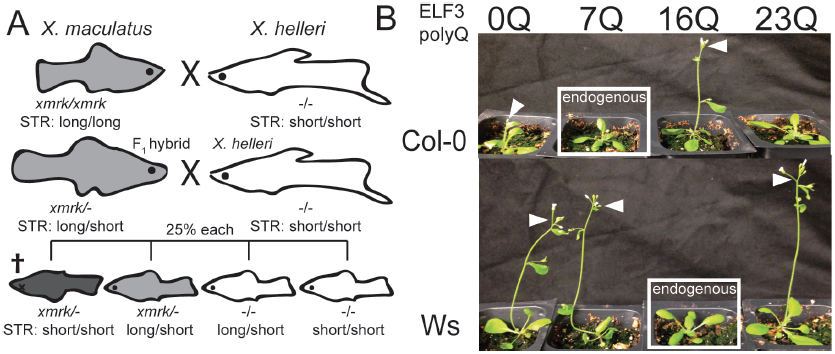
Genetic and transgenic analysis reveals STR-mediated incompatibilities. **A,** the Gordon-Kosswig-Anders cross shows a genetic incompatibility between two fish species in the *Xiphophorus* genus. Modified from Meierjohann and Schartl [63]. F_1_ hybrids back-crossed to their *X. helleri* parent yield a 3:1 ratio of viability, where the inviables result from co-segregation of the functional *xmrk* gene and a short STR allele in the *cdkn2a/b* promoter. Shading indicates melanism conferred by *xmrk*. **B,** genetic background is epistatic to effects of *ELF3* STR variation in *A. thaliana.* Expression-matched transgenic plants with various alleles of the *ELF3* STR in the Columbia (Col-0) and Wassilewskija (Ws) backgrounds, showing endogenous, exogenous, and synthetic (“0”) alleles in each background [25]. White boxes indicate transgenic plants carrying the *ELF3* STR endogenous to their respective background; white arrowheads indicate early-flowering ELF3 STR genotypes (*elf3* mutants and poorly-functioning *ELF3* STR alleles confer early flowering).

Similarly, we have shown that natural variation in the polyQ-encoding *ELF3* STR significantly affects all ELF3-dependent phenotypes in the plant *A. thaliana,* with *ELF3* STR length and phenotype showing a strikingly nonlinear relationship (Figure 1B)[25]. Some naturally occurring *ELF3* STR variants phenocopy *elf3*-loss-function mutants in a common reference background (Figure 1B), suggesting background-specific modifiers. Indeed, when we compare the phenotypic effects of each *ELF3* STR variant between two divergent backgrounds, Columbia (Col-0) and Wassilewskija (Ws), we find dramatic differences. The endogenous STR alleles from these two strains (Col-0 7 units, Ws 16 units) show mutual incompatibility when exchanged between backgrounds. The ELF3 protein is thought to function as an “adaptor protein” or physical bridge in diverse protein complexes [64,65]. We speculated that background-specific polymorphisms in these interacting proteins underlie the *ELF3* STR-dependent background effect.

Also in *A. thaliana*, a variable STR in the promoter of the *CONSTANS* gene has been linked to phenotypic variation in the onset of flowering [28]. *CONSTANS* encodes a major regulatory protein that promotes flowering. Transgenic experiments demonstrate that this regulatory STR variation affects *CONSTANS* expression and hence onset of flowering. However, the effects of this STR variation depend on the presence of a functional allele of *FRIGIDA*, a negative regulator of flowering that is highly polymorphic across *A. thaliana* populations.

A dramatic example of incompatibility can be found in an intronic repeat in the *IIL1* gene in *A. thaliana*, which was found to be dramatically expanded in one strain [55]. The expansion delayed flowering under high temperatures, but when crossed into the reference genetic background, a strongly interacting locus modifies this phenotype.

In the *Drosophila* genus, coding STR variation in the *per* gene co-evolves with other variants [59,66]. Transgenic flies expressing chimeric *per* genes with a *D. melanogaster* STR domain fused to a *D. pseudoobscura* flanking region (and vice versa) have arrhythmic circadian clocks, indicating the modifying effect of flanking variation in generating an STR-based genetic incompatibility. Among STRs subjected to exhaustive genetic study, to our knowledge, only the yeast *FLO1* coding STR has no known modifiers due to variation in genetic background [57].

In addition to these exhaustive genetic studies, there are several other observations that support the role of the genetic background in controlling the phenotypic effects of STRs. For instance, experiments in *Caenorhabditis elegans* and human cells indicate that the phenotypic effects of proteins with expanded polyQ tracts are modulated by genetic background [67], or by variants in interacting proteins [68]. In humans, genetic association studies indicate the existence of genetic modifiers of polyQ expansion disorders for both Huntington’s disease [69] and spinocerebellar ataxias [70]. Taken together, these experimental and observational data support our argument that functional STRs are likely to be enriched for variants in epistasis with other loci.

STRs with background-dependent phenotypic effects tend to either encode polyQ tracts or reside in promoter regions. There are good reasons to expect that these STR classes might be enriched in DNA/protein-protein interactions that could underlie epistasis. PolyQ tracts, specifically, often bind DNA surfaces [71], and an analysis of human protein interactome data found that polyQ-containing proteins engage in more physical interactions with other proteins than those without polyQs [72]. Similarly, noncoding STRs in regulatory regions may compensate for mutations in trans-acting factors, as observed for the STRs in the *cdkn2a/b* promoter in *Xiphophorus* [29] and in the *CONSTANS* promoter in *A. thaliana* [28]. We suggest that polymorphisms in protein interaction partners or in transcriptional regulators are plausible explanations for the observed background effects. In summary, we expect that STR variation is likely to contribute a substantial epistatic component to heritability, which has important implications for their use in explaining phenotypic variation.

## Analytical tools and genotyping methods continue to struggle with STR-specific challenges

To fulfill the promise of STR variation for explaining heritability, we need accurate, genome-wide assessment of STR variation in populations of humans and other organisms. The scientific community has tackled this problem in a flurry of recent studies describing methods for genotyping STRs genome-wide (Table 1). Specifically, in the last two years, several analytical tools have been developed to call STR genotypes from whole-genome-sequencing data [37–39]. These tools attempt to address the two major challenges for genotyping STRs: poor mappability due to low sequence complexity and high technical error rate due to amplification stutter.

**Table 1.** Technologies for assessing STR variation by high-throughput sequencing.

| Name | Data Source | Analysis strategy | Accepted coverage<sup>a</sup> | Reported accuracy | Reported efficiency | Limitations | Ref. |
| --- | --- | --- | --- | --- | --- | --- | --- |
| lobSTR | Human, whole-genome<sup>b,c</sup> | Align to modified reference | 1 read | 88%-95% | 0.2% of reads are informative | Depends on depth of sequencing and length of reads | [38] |
| RepeatSeq | Human, whole-genome<sup>b,d</sup> | Align to reference, locally realigned | 2 reads | 92% | Not reported | Depends on depth of sequencing and length of reads | [37] |
| STRViper | *A. thaliana* whole-genome<sup>b,d</sup> | Compare insert size to reference | 10 reads | 74% | Not reported | Cannot call STR unit number genotypes | [39] |
| Array Capture | Human, array capture<sup>b</sup> | RepeatSeq | 2 reads | 88%-92% | 2.2% informative reads | Low enrichment for STR-spanning reads | [35] |
| SureSelect RNA probe capture | Human, target enrichment, Roche 454 | Locally align flanking regions | 4 reads | 88%-95% | 27% informative reads | Expensive probe design, captured only 60% of targeted STRs | [36] |

a: Minimum coverage of a single STR that is considered sufficient to call a genotype.
b: Sequence data from Illumina HiSeq technology.
c: data references: [93,94]
d: data references: [95,96]
e: data references: [97,98]

To accurately map an STR sequence read and retrieve its unit number genotype, the sequence read must span the STR of interest and include some unique flanking sequence. This requirement limits the length of STRs that can be accurately genotyped and decreases effective STR coverage compared to average whole-genome-sequencing coverage (Figure 2). For this reason, much of the existing sequencing data, which consists largely of short reads (36 bp, 50 bp, or 76 bp) with only modest genome coverage (5-20X) is not suitable for accurate, genome-wide calls of STR genotypes; only a fraction of STRs, mostly short ones, can be assessed with some confidence (Figure 2).

**Figure 2.**
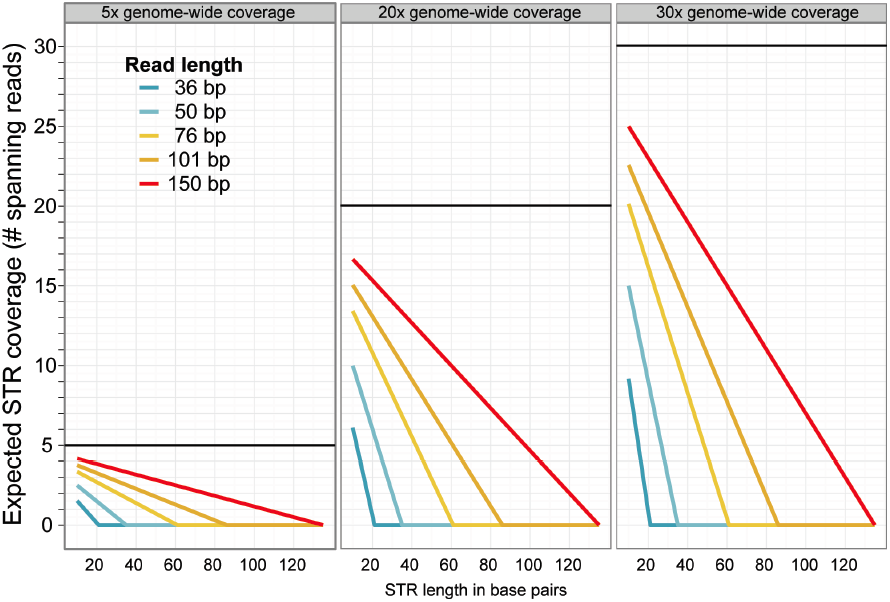
Effective reduction in STR coverage in whole-genome sequencing. Expected coverage of STRs for various sequencing depths and read lengths. We assumed 8 bp of flanking sequence on either side (per requirement for LobSTR software [38]). Black bars indicate nominal sequencing coverage for each scenario. 4-5X coverage (left panel) is typical for genomes in the human 1000 Genomes Project [95]; 15-20X coverage is typical for genomes in the *A. thaliana* 1001 Genomes Project [97,98].

Moreover, these analytical tools estimate technical error based on STR genotypes from sequenced homozygous or haploid genomes, ignoring somatic alleles within individuals (which are expected for STRs even in primary tissues, occurring at rates 10^4^-10^5^ times higher than SNV somatic mutations) [73–76]. Probabilistic error models have been formulated to quantify variation arising from technical sources [37,38], but in the face of somatic STR variation, these models presumably require substantial read coverage to call germ-line STR genotypes with confidence. However, because of the low effective coverage of STR loci (Figure 2), STR genotype calls are based on as few as one to two STR-spanning reads [37,38] (Table 1). Calls based on so few reads may not be accurate even for homozygous germline alleles. Calling heterozygous STR genotypes remains difficult with the modest coverage of most available whole-genome-sequencing data, such as found in the 1000 Genomes Project [12], which becomes even more challenging when potential somatic mutations contribute to a heterogeneous sample population. To illustrate this challenge, consider a heterozygous ∼30 bp-STR locus and whole-genome sequencing with 101 bp-reads at 5x coverage – this scenario is likely to yield just three STR-spanning reads (Figure 2). These three reads may represent one, two, or three different alleles, representing any mixture of two different germ-line alleles, somatic alleles, or technical error, making an accurate call difficult. Consequently, an increase in the sequencing depth of available data may be required before these tools reach their full potential.

Others have attempted to genotype STRs using whole-genome-sequencing data from paired-end reads (50bp) of size-selected genomic fragments [39], similar to strategies used to detect large insertions or deletions [77–80]. This approach is limited by the resolution of gel electrophoresis in the size selection of DNA fragments. Consequently, this method cannot determine STR unit number genotypes, but rather reports whether an STR is variable across samples. The authors argue that this approach is the most accurate for population-level detection of STR variability [81], but it is not informative for discerning the relationship between STR unit number genotype and phenotype.

Although these analysis tools represent important and useful advances, their limitations illustrate that ‘dustbin-diving’ of whole-genome-sequencing data may not suffice for accurate population-scale genotyping of STRs genome-wide. Alternative approaches that enrich for STR-spanning sequencing reads are needed. Indeed, two such approaches have been recently published. Both use targeted capture of STRs to enrich for STR-spanning reads combined with high-throughput sequencing compatible with midsize-reads (101 bp, 500 bp) [35,36]. Targeted STR capture requires the design of STR-specific probes (or rather probes specific to their unique flanking sequences) and involves additional sequencing, but these approaches can dramatically increase the number of informative reads, therefore providing substantial STR coverage for accurate genotyping calls (Table 1). For example, the SureSelect-RNA-probe capture method reports 27% informative STR-spanning reads compared to the 0.2 % informative reads found in whole-genome-sequencing data (Table 1). This increase in informative reads is a major advantage over whole-genome resequencing because STRs represent only a small fraction of the genome overall [35,36]. Although targeted capture combined with high-throughput sequencing appears to be a cost-effective alternative for accurate STR genotyping compared to whole-genome sequencing, distinguishing heterozygous alleles, somatic variants, and technical error remains a challenge. We suggest that recent innovations in single-molecule targeted capture [82] should be useful in distinguishing these categories and in further increasing enrichment of informative, STR-spanning reads.

## Lack of statistical models for detecting STR-phenotype associations in GWA

Assuming that we obtain accurate, population-scale genotype data for STRs, we may not yet have statistical tools appropriate for detecting STR associations with phenotype [8]. In diploid organisms, a biallelic SNV is typically analyzed by modeling phenotype as a function of the number of non-reference alleles at that locus (0, 1, or 2) in each individual. A null hypothesis of no monotonic relationship between phenotype and the allele count is then formulated and tested [83]. This framework cannot accommodate more than two alleles, which we would expect for many STRs. Simply using tagged SNVs linked to STRs to perform GWA is unfeasible, because linkage disequilibrium decays very quickly between SNVs and STRs across human populations [12].

To address these complications, a previous study attempted GWA between STR genotypes and human disease phenotypes by comparing relative frequencies of various alleles in pooled DNA from cases and controls [84]. By pooling samples, this approach eases the analysis of multiallelic loci, but it loses information by ignoring specific individuals.

In a more recent study, the authors used logistic regression and the analysis of variance to detect associations between STR alleles and quantitative phenotypes in an inbred *Drosophila* mapping population [13]. Given that significant associations were detected, such approaches may be sufficiently powerful in recombinant inbred lines. However, their strategy relied on homozygosity, and considered multiallelic STRs in a pairwise fashion, so these straightforward methods will lose power with outbred populations and multiallelic STRs.

The central confounder of these studies is that most STRs of appreciable variability (and thus, interest) are multiallelic, as a simple consequence of the STR mutational mechanism [17]. This multiallelic feature could be accommodated by treating STR alleles categorically, but this choice entails a corresponding reduction in power, because many alleles are rare.

Some studies have reported linear associations between STR unit number and quantitative phenotypes [27,57], suggesting that using simple tests of linear correlation between these variables may be a powerful option. However, this linearity (or even monotonicity) of the relationship between STR unit number genotype and phenotype is a poorly-supported assumption [25]. Nonetheless, STR unit number is a numerical variable, and it would be preferable to gain power from treating it as such. For instance, more similar STR unit number genotypes might be associated with more similar phenotypes, but this intuition may be difficult to generalize.

Lastly, both intuition (Box 1) and the studies discussed above lead us to expect that relatively many phenotypically relevant variable STRs will show epistasis with other loci. This epistasis will reduce power in tests of association between STRs and phenotype [85], given the inadequacy of the current paradigm of quantitative genetics in detecting and modeling the effects of epistasis [85,86]. At present, targeted and exhaustive genetic studies (as described above) are the only effective method for understanding the effects of epistasis.

In total, these obstacles present a daunting challenge for the integration of STR genotypes into the current genotype-phenotype maps. Overall, we call for a reappraisal of statistical methodologies for use in GWA with STR variation to account for these various STR-specific confounders.

## Somatic STR variation may be a sensitive marker for increased disease susceptibility

It has been appreciated for some time that the high STR mutation rate leads to somatic variation within individuals in addition to germ-line variation between individuals [71]. This somatic STR variation is particularly noticeable in tumor tissues, but is also measurable in primary tissues [73,87]. While these findings immediately led to systems of classification for tumor types and clones [76,88,89], the investigation of somatic STR variation (or MSI) may also inform us about general phenotypic states and disease susceptibility.

Patients with various complex diseases tend to carry a greater load of rare germ-line variants than unaffected control groups [6]. It is widely assumed that these rare variants contribute in some fashion to these disorders [90]; however, an alternative interpretation holds that they are signs of stochastic genome instability, which when increased leads to higher susceptibility to complex diseases. [6]. Increased genome instability will increase somatic variation, which may then serve as a read-out of disease susceptibility [6]. This alternative interpretation has some support from empirical data. For instance, perturbation of the molecular chaperone Hsp90, which stabilizes diverse DNA repair proteins, leads to increased somatic STR mutation rates in human cells; in various model organisms Hsp90 perturbation increases transposon mobility and intrachromosomal homologous recombination [31–34]. Hsp90 perturbation also increases the penetrance of many genetic variants in flies, plants, fish, worms and yeast, suggesting that increased genome instability and increased phenotypic heritability are associated [34]. If this association also applies to disease phenotypes, increased genome instability may predict higher disease susceptibility.

Consequently, although somatic MSI may not be the cause of disease phenotypes, it may serve as a biomarker for individuals who are more vulnerable to environmental and genetic perturbations leading to disease. Again, this strategy hinges on the development of cost-effective technologies for screening panels of STRs for somatic mutations across many humans, which will require new strategies to distinguish technical error from somatic STR variation.

Another possibility is that somatic variation is itself phenotypically relevant, or even plays a role in developmental processes. It is known that STRs are enriched in genes with neuronal function [91]; some have even proposed that such somatic mutation is a component of normal neuronal development in humans [92]. If this is the case, then a greater appreciation of somatic variation will be necessary to understand canonical developmental processes. Collectively, STR variation within (in addition to between) individuals has great potential as a read-out for disease susceptibility, and perhaps also as a cause of phenotypic variation itself.

## Concluding remarks

The study of STRs and other under-ascertained genomic elements has the potential to reshape our model of the heritability of complex diseases and traits, both in terms of the overall proportion of heritability explained, and in terms of the components of heritability themselves (Outstanding Questions). Experimental studies in model organisms have taught us that the phenotypic effects of genome-wide STR variation are both dramatic and impossible to understand without taking epistasis into account. In the future, our understanding will be improved by 1) accurate STR population-scale and somatic genotyping, 2) more appropriate statistical methods for analyzing STR-phenotype associations, and 3) a broader description of epistasis between STR variation and other loci in determining phenotype.

## OUTSTANDING QUESTIONS

- **In light of wide-spread epistasis, what statistical and experimental tools can quantify the effect of STR variation on phenotype?**
- **Can inexpensive, accurate tools be developed for germ-line and somatic STR genotyping?**
- **Will somatic STR variation be effective as a readout for disease susceptibility?**

## GLOSSARY

Short tandem repeat (STR): a repetitive nucleotide sequence that consists of many copies of a short sequence in tandem (ex. CAGCAGCAGCAG). STRs are frequently called **microsatellites**.
Single nucleotide variant (SNV): Variant that consists of a change at a single nucleotide position. Common SNVs are sometimes called single nucleotide polymorphisms (SNPs).
Heritability: The fraction of variation in a phenotype across a population that can be attributed to genetic differences.
Epistasis: Non-reciprocal interactions of non-allelic gene variants, due for instance to functional interdependence between gene products in a protein complex or metabolic pathway.
Genome-wide association (GWA): A set of methods by which each of a large number of genetic variants genome-wide is tested for statistical associations with a phenotype. Often referred to in the context of **genome-wide association studies** (**GWAS**).
Complex disease, complex traits: Complex diseases or traits are phenotypic characters thought to be affected by multiple genetic and environmental factors.
Somatic variation: Genetic variation across somatic cells or tissues of an organism, which are generally not inherited by offspring (which inherits instead **germ-line variation**). Generally arises from mutations in specific cell lineages after early development.
Microsatellite instability (MSI): Somatic variation of STRs (microsatellites) associated with phenotypic changes such as cancer, often due to mutations in DNA repair genes.
Bateson-Dobzhansky-Muller incompatibility: Hybrid incompatibilities observed when crossing two close species or divergent strains of a species with one another. Caused by the co-segregation of non-parental allele combinations, resulting in a dysfunctional genetic interaction (negative epistasis).
Genetic anticipation: A mode of disease inheritance characterized by progressively earlier ages of disease onset as generations progress. Generally caused by the gradual expansion of STRs.

## Acknowledgements

This work was supported by grants from the National Human Genome Research Institute Interdisciplinary Training in Genomic Sciences (2T32HG35-16 to MOP, T32 HG00035 to KDC) and the National Institute of Health New Innovator Award (DP2OD008371 to CQ). We are grateful to members of the Queitsch and Shendure laboratories for helpful discussions, and to Wen Huang and David Mittelman for responding to email inquiries.

